# Female-biased expressed odorant receptor genes differentially tuned to repulsive or attractive plant volatile compounds in the turnip moths

**DOI:** 10.1101/2023.07.11.548602

**Authors:** Dan-Dan Zhang, Xiao-Qing Hou, Daniel Powell, Christer Löfstedt

## Abstract

Insects rely on their highly efficient and precise olfactory systems to find suitable mates, host plants and oviposition sites, and adapt to the changing environment. The odorant receptors (ORs) including pheromone receptors (PRs) play a vital role in this process. While extensive studies have been focusing on deorphanization of lepidopteran PR genes, the information on the ligand profiles of general ORs is still sparse. In the present study, we identified a repertoire of 61 ORs including the co-receptor Orco from antennal and ovipositor transcriptomes of the turnip moth *Agrotis segetum*, which clustered in all the major lepidopteran OR clades. We characterized the function of eight female-biased expressed ORs in *Xenopus* oocytes and found three ORs differentially tuned to plant volatile compounds that might be repulsive or attractive to the moths. AsegOR13 was broadly tuned to a number of herbivore-induced plant volatiles (HIPVs) while AsegOR20 was specific to citral; AsegOR17 was narrowly tuned to the alcohols, isoamyl alcohol, pentanol and benzyl alcohol, that are potentially attractive to moths. The orthologues of the three ORs in other moth species seem to share the conserved function. Our results support the hypothesis that insects recognize their host plants mostly by detecting the mixture of ubiquitous compounds, instead of taxonomically characteristic host compounds. The combination of narrowly and broadly tuned ORs will ensure both the accuracy of the most important odor signals and the plasticity of the olfactory system to the changes in the environment.

## Introduction

Insects have succeeded to survive and thrive in the world filled with complex odors. This is owed to their highly efficient and precise olfactory systems, which guide them to find suitable mates, host plants and oviposition sites, and adapt to the ever-changing environment (Bruce et al., 2005; Bruyne & Baker, 2008; Renou, 2014). Olfactory receptors equipped on the antennae of insects play an essential role in this process, detecting various odorants and converting the chemical signals to electrical signals that are further transferred to the brain (Leal, 2013). Insect olfactory receptors are encoded by two major gene families, odorant receptor genes (ORs) and ionotropic receptor genes (IRs) (Silbering et al., 2011; Wicher & Miazzi, 2021; Hou et al., 2022). While IRs are more ancient and primarily respond to acids and amines, ORs evolved in parallel with the onset of insect flight and sensitively detect pheromones and a range of plant volatiles (Wicher, 2018 and therein).

Insect ORs are heteromeric complexes, each consisting of a highly conserved olfactory co-receptor (Orco) subunit and a more divergent tuning OR subunit that accounts for the variable response spectra (Sato et al., 2008). There have been extensive studies on the ORs detecting female-released sex pheromones, the so-called pheromone receptors (PRs), in moths that are well-known for their selective and sensitive sex pheromone communication systems (Zhang & Löfstedt, 2015). The majority of the PRs characterized so far are responsive to Type I pheromones (fatty alcohols, aldehydes or fatty alcohol acetates) (Löfstedt et al., 2016); mostly clustered in the classical PR clade (Zhang & Löfstedt, 2015), with some fall into the so-called novel PR clade (Bastin-Héline et al., 2019; Yuvaraj et al., 2022). It seems that the PRs for the Type II epoxide pheromones and the ancestral Type 0 pheromones (short-chain secondary alcohols or ketones) also cluster in respective clades (Li et al., 2017; Yuvaraj et al., 2017), while the receptors for Type II hydrocarbon pheromones reside in classical PR clade (Zhang et al., 2016). PRs have typical male-biased expression pattern, either exclusively expressed in male antennae, or significantly more abundant in male antennae compared to female antennae (Zhang & Löfstedt, 2015).

Much less is known about the response spectra of ORs other than PRs in moths. Evidence from previous studies suggested that many of these ORs recognize plant volatile compounds and might play important roles in insect-plant interaction (Tanaka et al., 2009; de Fouchier et al., 2017; Guo et al., 2021). In the present study, we identified a full OR repertoire from the transcriptome of the turnip moth, *Agrotis segetum*, and characterized the functions of eight ORs that were relatively abundant and female-biased expressed. We found three of these ORs had different response spectra to plant volatiles that are potentially attractive or repulsive to the herbivorous moths.

## Materials and Methods

### Chemicals

A panel of 48 odorants, consisting of the common plant volatiles and pheromone related compounds (Additional file 1: Table S1), were tested against AsegORs in *Xenopus* oocyte recordings. The plant volatile compounds included different chemical classes: monoterpenes/monoterpenoids, esters, aldehydes and alcohols. The stock solutions of 100 mM were prepared in dimethyl sulfoxide (DMSO) and stored at −20 °C. The working solutions with indicated concentrations were diluted from the stock solutions by Ringer’s buffer (96 mM NaCl, 2 mM KCl, 5 mM MgCl2, 0.8 mM CaCl2, 5 mM HEPES, pH 7.6) before each experiment. Each experiment included a negative control with Ringer’s buffer containing 0.1% DMSO.

### Insects

The *A. segetum* moths were from the lab culture at the Pheromone Group, Department of Biology, Lund University. Larvae were reared on artificial bean-based diet in a climate chamber at 25°C with 50% of relative humidity and a light/dark cycle of 16:8. The antennae were dissected from 3 days old male virgin moths that were separated from the females since pupal stage and were fed with 10% honey solution after emergence. The collected antennae in the Eppendorf tubes were immediately frozen on dry ice and stored at – 80 °C until further use.

### Gene annotation and expression level estimation

Functional annotation of gene assemblies was performed by blasting against the pooled database of nr (NCBI non-redundant protein sequences), KOG/COG (Clusters of Orthologous Groups of proteins) and Swiss-Prot with a cut-off E-value of 1e−5. With the identified AsegOR sequences as queries and the transcriptome as a custom database, additional blast searches were performed to ensure that all OR genes were discovered. The expression levels of AsegORs were normalized across sequencing libraries using the scaling factor transcripts per million (TPM) (Li et al., 2011).

### Phylogenetic analysis

The phylogenetic analysis with the amino acid sequences of ORs from *A. segetum, Helicoverpa armigera* (Liu et al., 2012), *Bombyx mori* (Wanner et al., 2007), *Spodoptera littoralis* (Poivet et al., 2013) and *Epiphyas postvittana* (with genes in the novel PR clade) (Corcoran et al., 2015), *Ectropis grisescens* (Li et al., 2017) and *Operophtera brumata* (Zhang et al., 2016) (with receptors for Type II sex pheromones) and *Eriocrania semipurpurella* (with receptors for Type 0 sex pheromones) (Yuvaraj et al., 2018). The sequences with ≥ 200 amino acids were aligned with MAFFT. The maximum-likelihood tree was constructed by IQ-TREE with the WAG+G+F substitution model decided by “Find best protein model”; bootstrap analysis of 100 replicates was performed. The tree was rooted with the Orco lineage and further edited in FigTree 1.4.4.

### Molecular cloning

The antennae were homogenized with plastic pestles in lysis buffer RLT Plus on ice. The total RNAs were extracted using RNeasy Plus Micro Kit (Qiagen, GmbH, Hilden, Germany) and reverse transcribed into cDNAs with the SuperScript™ IV First-Strand Synthesis System (Invitrogen, Carlsbad, CA, USA) following the manufacturer’s instruction. The full-length of OR genes were amplified with primers containing Kozak sequence (“GCC ACC”) and restriction sites (Additional file 1: Table S2) using Platinum Pfu Polymerase (Thermo Fisher Scientific), and then sub-cloned into pCS2+ expression vectors. Sanger sequencing was performed to verify the gene sequences, using a capillary 3130xL Genetic Analyzer (Thermo Fisher Scientific, Waltham, MA, USA) at the sequencing facility at Department of Biology. The recombinant pCS2+ plasmids containing the ORs were purified using the PureLinkTM HiPure Plasmid Filter Midiprep Kit (Thermo Fisher Scientific). The plasmids were linearized by NotI (Promega) and transcribed to cRNAs using mMESSAGE mMACHINE SP6 transcription kit (Thermo Fisher Scientific).

### Functional characterization of ORs in Xenopus oocytes

The heterologous expression of ORs in *Xenopus* oocytes and two-electrode voltage clamp recordings were performed following previously described protocols (Hou et al., 2020). In brief, oocytes were collected from the ovary of female *X. laevis* (supplied by Centre d’Elevage de Xénopes, Rennes, France) and treated with 1.5 mg/mL of collagenase (Sigma-Aldrich Co., St. Louis, MO, USA) dissolved in Ca^2+^-free Oocyte Ringer 2 buffer (82.5 mM NaCl, 2 mM KCl, 1 mM MgCl2, 5 mM HEPES, pH 7.5) at room temperature for 15–18 min. Mature and healthy oocytes (stage V–VII) were isolated and co-injected with the cRNAs of each AsegOR and the co-receptor AsegOrco (50 ng each), and incubated for 4–7 days at 18 °C in Ringer’s buffer containing 550 mg/L sodium pyruvate and 100 mg/L gentamicin. The ligand-induced whole-cell inward currents were recorded with a TEC-03BF signal amplifier at a holding potential of −80 mV. The solutions were delivered to the oocytes through a computer-controlled perfusion system with compound stimuli at a rate of 2 mL/min for 20 s and extensive washing in Ringer’s buffer at a rate of 4 mL/min between stimulations. Data collection and analysis were carried out by Cellworks software (npi electronic GmbH,Tamm, Germany).

## Results

### Identification of the A. segetum OR repertoire from antennal and ovipositor transcriptomes

We identified the OR gene repertoire from the transcriptomes of the antennae (N=4 for both sexes) and ovipositor (N=2) that we recently sequenced (Bioproject accession number PRJNA707654; Hou et al., 2022). We identified 61 full-length OR sequences in total, including Orco and the PRs we previously reported (Zhang & Löfstedt, 2013).

The PRs that were functionally characterized in our previous study had been named as AsegOR1 and AsegOR3-10 (Zhang & Löfstedt, 2013). “AsegOR2” was not used to avoid being confused with Orco which was formerly named as OR2 in other moth species. We identified two additional PR candidate genes from the transcriptome hereby and named them as AsegOR11 and AsegOR12. The eight female-biased expressed ORs that we functionally characterized in this study were named as AsegOR13-20. Thereafter, the other candidate ORs were named as AsegOR21-59 according to the order of their contig numbers in the transcriptome. Some genes have more than one copy and were hence name as AsegOR23a/b/c and AsegOR40a/b.

### Clustering of A. segetum ORs with the major lepidopteran OR clades

Lepidopteran ORs were classified into 23 major clades (A-W) in previous studies (de Fouchier et al., 2017; Guo et al., 2021). The AsegORs that we identified clustered into these major clades (Figure 1), with AsegOrco in clade A (the Orco clade) and the PRs (AsegOR1 and AsegOR3-10) in clade V (the classic PR clade). Intriguingly, we identified another candidate gene (AsegOR11) in the clade V, as well as candidate genes (AsegOR12, AsegOR33 and AsegOR54) in clade U (the so-called novel PR clade) (Bastin-Heline et al., 2019; Yuvaraj et al., 2022). The eight female-biased expressed ORs that we investigated in this study were scattered in the phylogenetic tree, instead of clustering together as the male-biased expressed PR genes.

**Figure 1.**
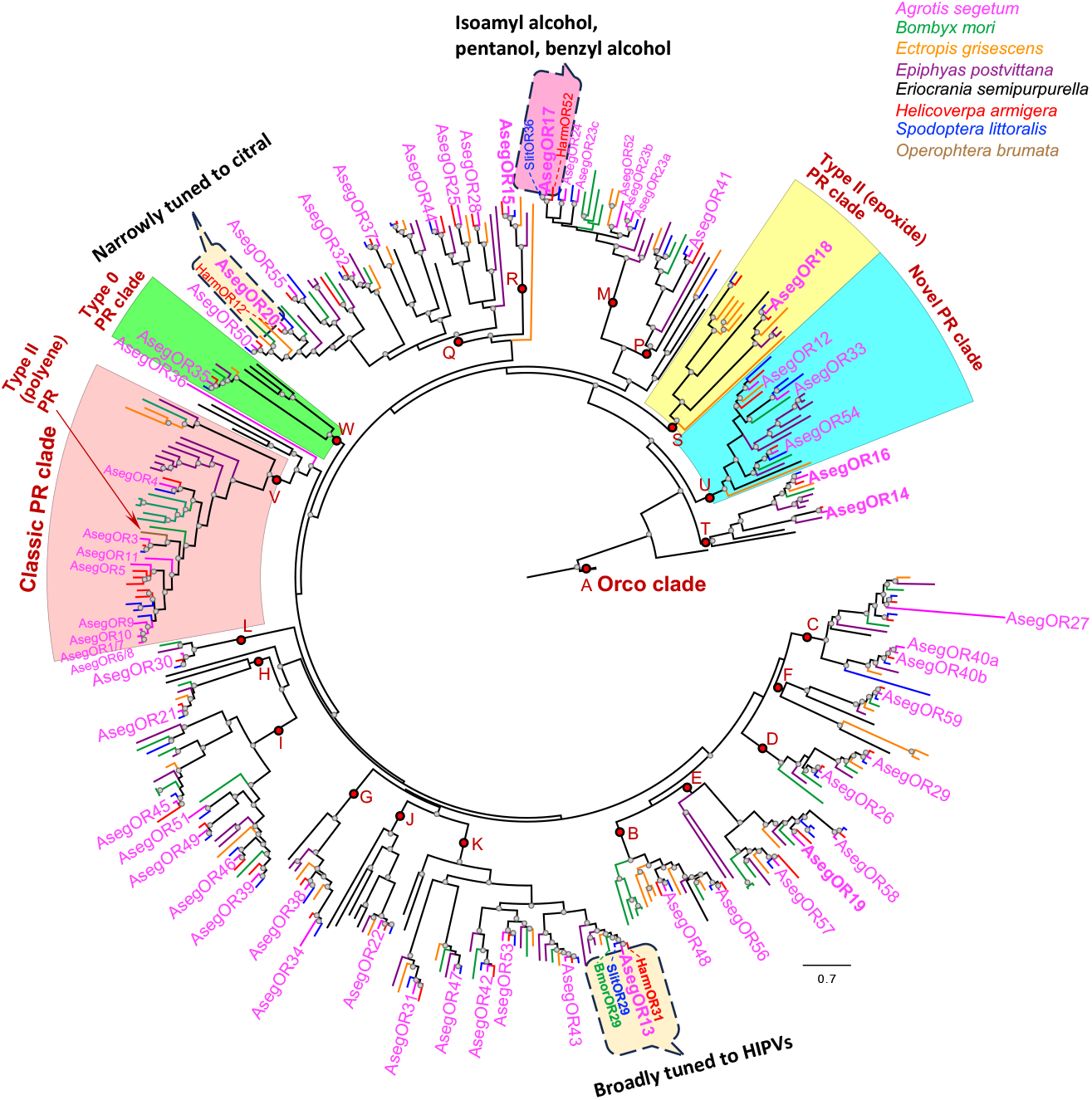
Phylogeny of Lepidoptera ORs. The maximum-likelihood phylogenetic tree was based on the amino acid sequences (≥ 200 aa) from *A. segetum, H. armigera, B. mori, S. littoralis, E. postvittana, E. grisescens, O. brumata* and *E. semipurpurella*. The tree was rooted with Orco lineage; bootstrap values (N = 100) over 70 were shown on the branches with small grey circles. The branches were color coded based on species as indicated in the legend. The AsegOR gene names were labeled on the tips. The eight female-biased expressed ORs that we characterized in this study were highlighted with bold font. The major lepidopteran OR clades (A-W, following Guo et al., 2021) were shown with red circles on the branches leading to the clades, with Orco clade, classic PR clade, novel PR clade, Type 0 PR clade and Type II (epoxide or polyene) PR (clade) noted at respective positions. The deorphanized AsegOR13, AsegOR17 and AsegOR20 and their orthologues in other species were framed in dashed line dialog boxes (pink for attractants and yellow for repulsive HIPVs) with the ligand profile information beside. The ligand profiles of the other species were from previous studies (Anderson et al., 2009; Tanaka et al., 2009; De Fouchier et al., 2017; Di et al., 2017; Guo et al., 2021).

### The expression pattern of AsegORs

The AsegORs were predominately expressed in the antennae, except for a few that were lowly expressed in the ovipositors (*e*.*g*., AsegOR12, AsegOR21, AsegOR22 and AsegOR44). The co-receptor, AsegOrco, was arguably the most abundant OR in the antennae of both males (TPM = 656.21 ± 39.80, N = 4) and females (TPM = 309.79 ± 10.65, N = 4) (Figure 2). Genes in the PR clade had the highest expression levels among the tuning ORs (Figure 2). AsegOR9, the PR that were specific to the major pheromone component (*Z*)-5-decenyl acetate (*Z*5-10:OAc) (Zhang & Löfstedt, 2013), was enriched in male antennae (TPM = 435.45 ± 31.45, N = 4) whereas not expressed in female antennae (TPM = 0 in all four replicates). AsegOR4 and AsegOR5, specific receptors for the other pheromone components (*Z*)-7-dodecenyl acetate (*Z*7-12:OAc) and (*Z*)-9-tetradecenyl acetate (*Z*9-14:OAc) respectively, also showed typical male-biased expression, with female to male expression (F/M ratio) as 6.02E-4 and 0.021. The paralogous genes AsegOR1/7, AsegOR6/8 and AsegOR10 that we previously identified shared high sequence identities with each other (82.9% - 98.4%) and were presented in a large polycistronic transcript with significantly higher expression level in male antennae (190.62 ± 11.85, N = 4) compared to female antennae (4.83 ± 0.79, N = 4). The new candidate gene we identified in the PR clade (AsegOR11) and the gene in the novel PR clade (AsegOR12) were male-biased expressed as well (F/M ratio = 4.08E-3 and 0.15 respectively), suggesting their potential roles in detecting female-produced sex pheromone components. AsegOR3, however, was not male-biased expressed although it located in the PR clade (F/M ratio = 1.08). It was in the previously “ligand unknown” Cluster III (Krieger et al., 2009; Baker, 2009; Zhang & Löfstedt, 2013) and was shown as responding to polyenic hydrocarbons (Zhang et al., 2016). The other two genes in the novel PR clade (AsegOR33 and AsegOR54) showed female-biased expression, with a F/M ratio of 15.3 and 1.95 respectively.

**Figure 2.**
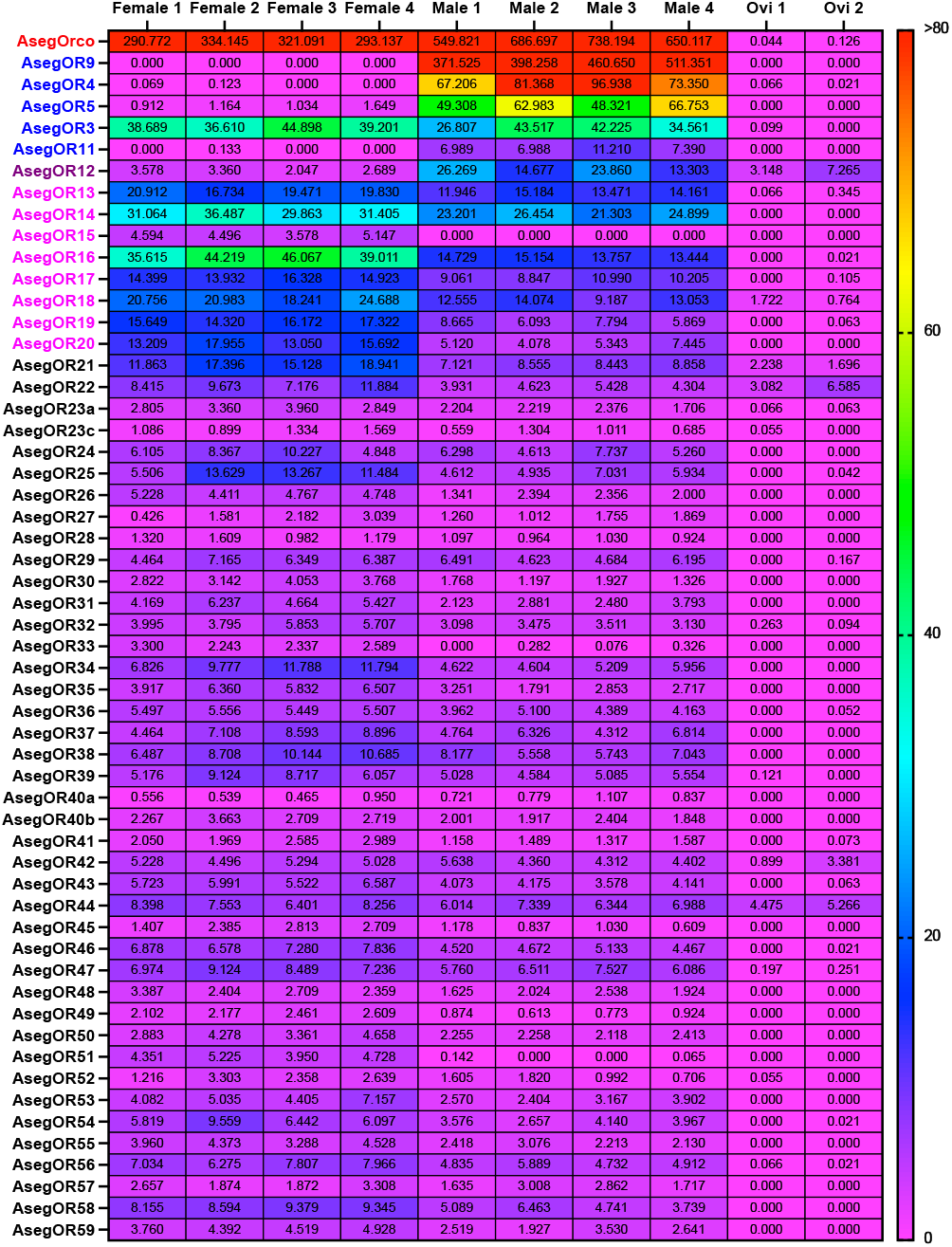
The expression profiles of AsegORs. The expression levels of AsegORs in different female antennae (Female 1-4) and male antennae (Male 1-4) samples as well as ovipositors (Ovi 1/2) were illustrated in a heatmap with TPM values (transcripts per million).

The eight ORs investigated hereby (AsegOR13-AsegOR20), on the contrast, had female-biased expression pattern, including one (AsegOR15) exclusively expressed in female antennae and the others with F/M ratio ranging from 1.34 to 2.89 (Figure 2). They all had reasonably high expression levels in female antennae (TPM = 14.90 ± 0.52 to 41.23 ± 2.39, N = 4) except for AsegOR15 the TPM of which was 4.45 ± 0.33. All the other ORs were lowly expressed with TPM ≤ 10 in both female and male antennae (Figure 2).

### The tuning profiles of female-biased expressed ORs

We next examined the tuning profiles of the eight ORs (AsegOR13-AsegOR20) that were abundant and female-biased expressed. Each of the eight genes was co-expressed with AsegOrco in *Xenopus* oocytes and tested against a panel of 48 compounds including common plant volatile compounds and the pheromone related compounds. Of the eight ORs, five did not show significant response to all tested compounds and three were responsive to different plant volatiles whereas no response to pheromone related compounds.

Both AsegOR13 and AsegOR20 responded to monoterpenes and monoterpenoids at the concentration of 100 μM but with different tuning breadths. AsegOR13 was broadly tuned, with the most potent ligands as citral and myrcene, followed by geraniol, linalool, (-)-limonene and (-)-β-pinene, as well as an ester (*Z*)-3-hexenyl acetate (Figure 3A and 3B). Citral and myrcene also exhibited higher sensitivities compared to other ligands in dose response trials (Figure 3C). AsegOR20 was specifically tuned to citral (Figure 4A and 4B) and the response was dose dependent (Figure 4C); it also had minor responses to geraniol and linalool (Figure 4A and 4B).

**Figure 3.**
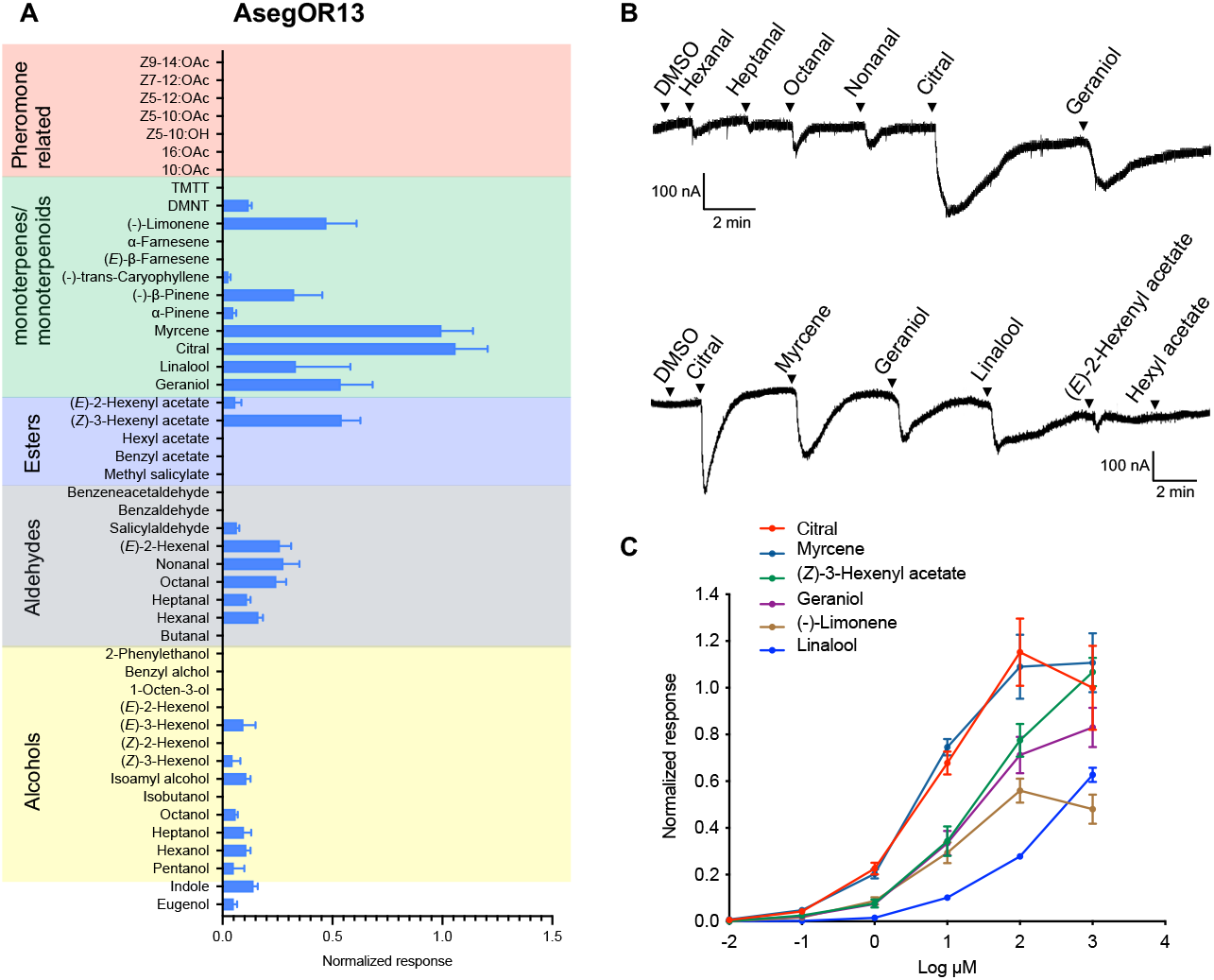
AsegOR13 was broadly tuned to monoterpenes/monoterpenoids in *Xenopus* oocytes. **(A)** The mean values ± SE of the currents in nano-Ampere (nA) when stimulated by compounds at 100 μM, N=3-13. **(B)** Representative current traces of oocytes co-injected with cRNAs encoding AsegOR13 and AsegOrco upon successive exposures to 100 μM stimuli. Each compound was applied for 10 s at the time point indicated by arrows. **(C)** The responses of AsegOR13/Orco to different doses of each compound, N=3-5.

**Figure 4.**
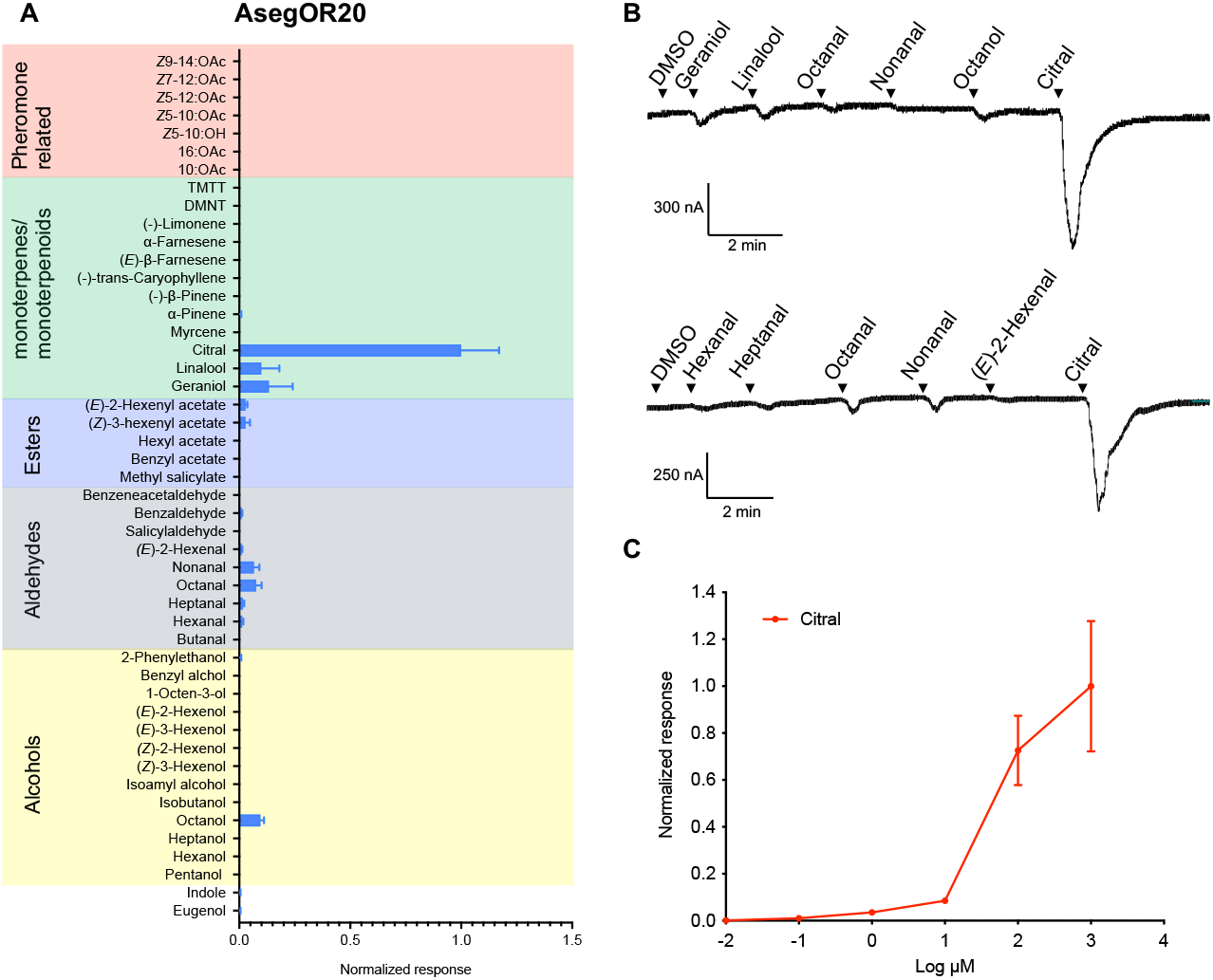
AsegOR20 was narrowly tuned to citral in *Xenopus* oocytes. **(A)** The mean values ± SE of the currents in nano-Ampere (nA) when stimulated by compounds at 100 μM, N=3-10. **(B)** Representative current traces of oocytes co-injected with cRNAs encoding AsegOR20 and AsegOrco upon successive exposures to 100 μM stimuli. Each compound was applied for 10 s at the time point indicated by arrows. **(C)** The responses of AsegOR20/Orco to different doses of each compound, N=3-5.

AsegOR17 was narrowly tuned to alcohols, with the largest response to isoamyl alcohol and smaller responses to pentanol and benzyl alcohol (Figure 5A and 5B). The responses to the active ligands were dose dependent (Figure 5C).

**Figure 5.**
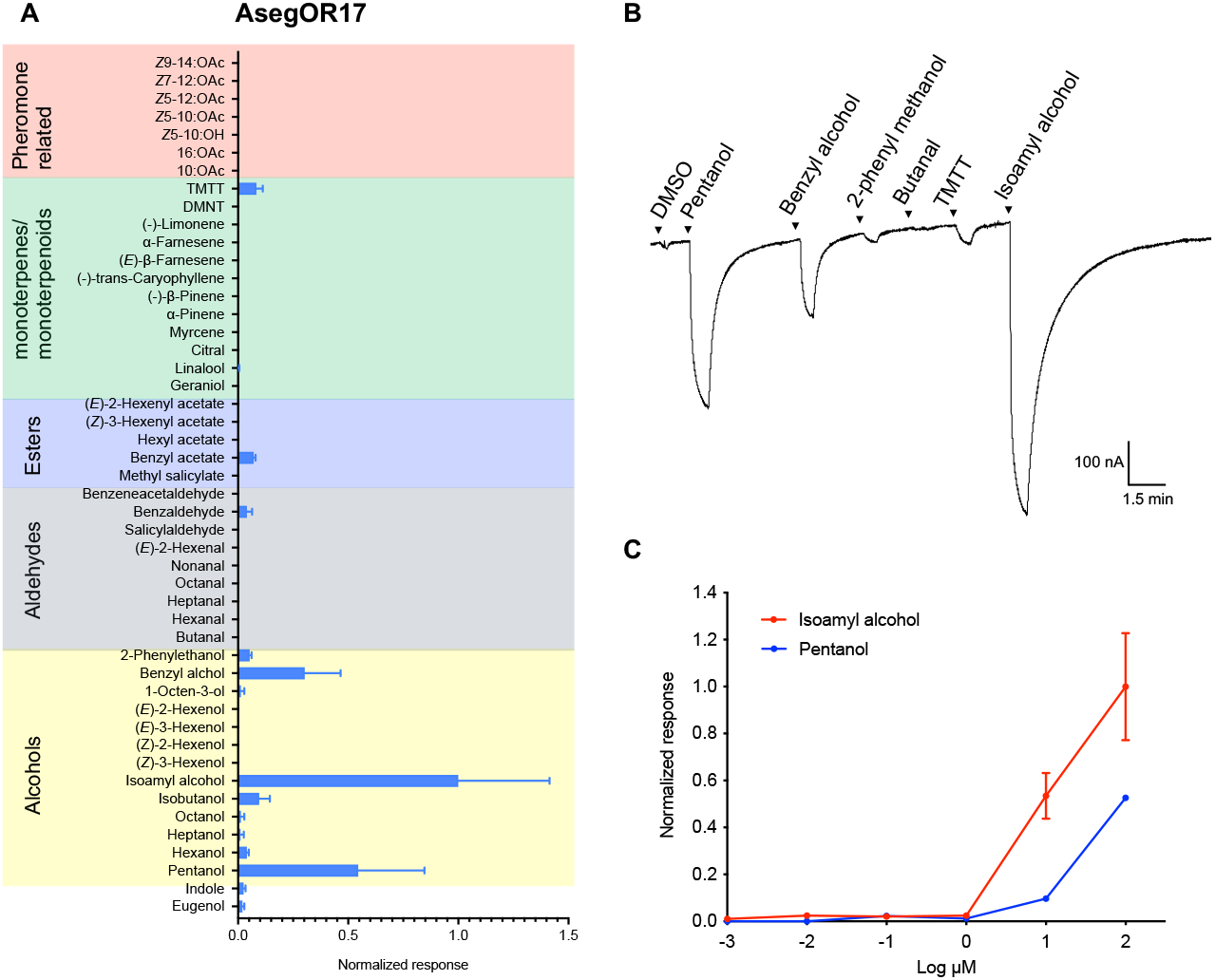
AsegOR17 was narrowly tuned to isoamyl alcohol, pentanol and benzyl alcohol in *Xenopus* oocytes. **(A)** The mean values ± SE of the currents in nano-Ampere (nA) when stimulated by compounds at 100 μM, N=3-9. **(B)** Representative current traces of an oocyte co-injected with cRNAs encoding AsegOR17 and AsegOrco upon successive exposures to 100 μM stimuli. Each compound was applied for 10 s at the time point indicated by arrows. **(C)** The responses of AsegOR17/Orco to different doses of each compound, N=3-5.

## Discussion

It is a vital and challenging task for the herbivore insects to locate the host plants and avoid unsuitable hosts. Insects fulfill this task thanks to their highly efficient olfactory systems, which enable them to recognize relevant host plant volatiles among the fast changing and complex background. Plant volatiles are often a blend of several hundred compounds (Bruce et al., 2005). Insects mostly rely on the mixture of ubiquitous compounds, instead of the “signature compounds” for specific plant species or families, to find their hosts; the ratio of the ubiquitous compounds released by the plants is important information for the insects to determine the identity and status of the plants (Bruce & Pickett, 2011).

In this study, we identified the ORs of the *A. segetum* moths in all the major lepidopteran OR clades, indicating that the species should have the capacity to recognize ubiquitous plant volatile compounds. This is in line with their generalist lifestyle, feeding on a wide range of plants from different families. Nevertheless, moth species with different host plant preferences and even the specialists (*e*.*g. B. mori* that only feed on the leaves of mulberry) were also well-represented in most of the OR clades. This suggested that lepidopteran ORs are at least genetically conserved; the generalists and specialists have similar peripheral olfactory systems, while the various host preferences might be due to the processing of information in central nervous systems (CNS), consistent with previous *in vivo* observations (Bruce & Pickett, 2011). Such an arrangement provides flexibility to adapt to the changing environment and possibility to switch the hosts.

Whether or not the conserved OR repertoire also have conserved response profiles remains elusive. Previous studies found some OR orthologues that were functionally conserved across taxa, *e*.*g*. the OR lineage specific to the key floral odorant phenylacetaldehyde (PAA) that were conserved across moths and butterflies (Guo et al., 2021). However, it is not possible to draw a firm conclusion on whether this functional conservation is general for all the OR clades, as studies of comprehensive deorphanization of ORs are still sparse and the data were collected from different expression systems, with different odorant panels and concentrations. We found that AsegOR13 was broadly tuned to monoterpenes/monoterpenoids, including citral, myrcene, limonene, linalool, geraniol, β-pinene and DMNT, and also to (*Z*)- 3-hexenyl acetate. These are typical herbivore-induced plant volatiles (HIPVs) that have been shown to either repel the herbivorous moths or attract their parasitoids, thus acting as direct or indirect defense approach for the plants against herbivory (Dicke & Baldwin, 2010; Gish et al., 2015; Guo & Wang, 2019). It seems that its orthologues in clade K share the conserved function as receptors with broad spectra to HIPVs, although not having exactly the same ligand profiles due to different test panels: SlitOR29 in *S. littoralis* broadly tuned to terpenes/terpenoids including ocimene, myrcene, carene, geraniol, DMNT and sulcatone, as well as (*Z*)-3-hexenyl acetate (de Fouchier et al., 2017); HarmOR31 in *H. armigera* broadly tuned to myrcene, geraniol, (*Z*)-3-hexenyl acetate, citral, 1-octanol, *cis*-jasmone, etc (Di et al., 2017; Guo et al., 2021); BmorOR29 in *B. mori* to citral, linalool, etc (Tanaka et al., 2009).

AsegOR20 was specific to the HIPV citral. Its orthologue HarmOR12 in clade Q was also narrowly tuned to citral (Di et al., 2017), supporting the functional conservation in this clade. The other orthologues in the same clade, SlitOR12 in *S. littoralis* and SlituOR12 in *S. litura* were not tested against citral (Zhang et al., 2013; de Fouchier et al., 2017). It is therefore unknown how general the functional conservation is across species.

AsegOR17 was narrowly tuned to isoamyl alcohol (also known as 3-methyl-1-butanol), pentanol and benzyl alcohol. The orthologue HarmOR52 in clade M was also responsive to these three alcohols (Di et al., 2017; Guo et al., 2021); SlitOR36 responded to several alcohols with benzyl alcohol as the best ligand, while isoamyl alcohol and pentanol were not tested (de Fouchier, 2017). Isoamyl alcohol, the best ligand for AsegOR17, naturally occurs in fermenting molasses and is produced by yeast (Utrio & Eriksson 1977; Vitanović et al., 2020). It has been shown to be attractive to female moths (including Noctuidae, Gelechiidae and Erebidae), flies and mosquitos (Landolt & Hammond, 2001; Tóth et al., 2010; Verhulst et al., 2010, 2011; Vitanović et al., 2020). The secondary ligands pentanol and benzyl alcohol were also reported as attractants for moths (Sun et al., 2014; Di et al., 2017; de Fouchier et al., 2018).

Among the three AsegORs that we deorphanized, AsegOR17 and AsegOR20 were narrowly tuned to the ubiquitous plant volatile compounds, supporting that general ORs can be as specific as PRs. The narrowly tuned ORs are often involved in labelled line olfactory circuits, associated with the chemical cues that are important for insects’ survival and reproduction (Haverkamp et al., 2018; Guo et al., 2021; Wicher & Miazzi, 2021). Of the best ligands for these two ORs, isoamyl alcohol was attractive to moths while citral was repulsive. They might provide information about the status of the plants, such as the sufficiency of molasses as food sources and the degree of infestation, therefore playing important roles for the moths to assess the suitability of the plants as hosts. It is worthwhile testing whether the specific responses to isoamyl alcohol and citral are well-conserved in the orthologous ORs across the taxa.

Meanwhile, we also found an OR that was broadly tuned to the repulsive HIPVs. This is reminiscent of some PRs broadly tuned to behavioral antagonists (Zhang & Löfstedt, 2015; Zhang, 2021). In both cases, it is conceivable that the activation of a general neuron for repulsive compounds is sufficient to abort the flight of moths towards the source (Takanashi et al., 2006). The broadly tuned olfactory receptors were considered to be associated with combinatorial coding of the olfactory cues, and they could provide the adaptive advantage to exploit a wide range of odors in the environment and track the changes in the odor signals (Andersson et al., 2015; Haverkamp et al., 2018; Wicher, 2018). The combination of narrowly and broadly tuned ORs will ensure both the accuracy of the most important odor signals and the plasticity of the olfactory system to the changing environment.

## Author Contributions

X.Q.H, C.L. and D.D.Z. designed the research. X.Q.H and D.D.Z. performed the research. C.L. and D.D.Z. contributed to project supervision, as well as new reagents and analytical tools. X.Q.H, D.P. and D.D.Z. analyzed the data. D.D.Z. wrote the paper with inputs from all the co-authors.

## Notes

### Competing Interest Statement

The authors have declared no competing interest.

