## Additional file 1 for "Female-biased expressed odorant receptor genes differentially tuned to repulsive or attractive plant volatile compounds in the turnip moths"

**Table S1.** Compounds used for characterization of *Agrotis segetum* odorant receptors (ORs), including their purities and source information.

| Class | Compounds | Purity (%) | Source |
|---|---|---|---|
| Pheromone related compounds | *Z*9-14:OAc | >96 | Lab stock |
| | *Z*7-12:OAc | >96 | Lab stock |
| | *Z*5-12:OAc | >96 | Lab stock |
| | *Z*5-10:OAc | >96 | Lab stock |
| | *Z*5-10:OH | >96 | Lab stock |
| | 16:OAc | >96 | Lab stock |
| | 10:OAc | >96 | Lab stock |
| Monoterpenes/monoterpenoids | TMTT | -/- | trc |
| | DMNT | -/- | trc |
| | (-)-limonene | 96 | Sigma-Aldrich |
| | α-farnesene | 99 | Sigma-Aldrich |
| | (*E*)-ß-farnesene | >90 | Aldrich |
| | (-)-*trans*-caryophyllene | >98.5 | Sigma |
| | (-)-ß-pinene | 99 | Aldrich |
| | α-pinene | 98 | Sigma-Aldrich |
| | myrcene | >90 | Sigma-Aldrich |
| | citral | 95 | Aldrich |
| | linalool | ≥97 | Aldrich |
| | geraniol | 98 | Aldrich |
| Esters | (*E*)-2-hexenyl acetate | 97 | Aldrich |
| | (*Z*)-3-hexenyl acetate | 96 | Sigma-Aldrich |
| | hexyl acetate | 99 | Aldrich |
| | benzyl acetate | ≥99 | Sigma-Aldrich |
| | methyl salicylate | 99 | Sigma-Aldrich |
| Aldehydes | benzeneacetaldehyde | 97.5 | Sigma-Aldrich |
| | benzaldehyde | 99.5 | Aldrich |
| | salicylaldehyde | 98 | Sigma-Aldrich |
| | (*E*)-2-hexenal | 98 | Aldrich |
| | nonanal | >95 | Sigma-Aldrich |
| | octanal | 99 | Aldrich |
| | heptanal | 95 | Aldrich |
| | hexanal | 98 | Aldrich |
| | butanal | 99.5 | Aldrich |
| Alcohols | phenylethyl alcohol | 99 | Aldrich |
| | benzyl alcohol | 99 | Sigma-Aldrich |
| | 1-octen-3-ol | 98 | Aldrich |
| | (*E*)-2-hexenol | 96 | Aldrich |
| | (*E*)-3-hexenol | 97 | Aldrich |
| | (*Z*)-2-hexenol | 95 | Aldrich |
| | (*Z*)-3-hexenol | 98 | Aldrich |
| | isoamyl alcohol | ≥98.5 | Sigma |
| | isobutanol | ≥98.5 | Sigma |
| | octanol | ≥99 | Sigma-Aldrich |
| | heptanol | 98 | Aldrich |
| | hexanol | 98 | Aldrich |
| | pentanol | ≥99 | Sigma-Aldrich |
| Other | indole | 99 | Aldrich |
| | eugenol | 99 | Aldrich |

trc: Toronto Research Chemicals.

**Table S2.** Primers used in this study.

| Genes | Primer sequence (5'-3') |
|---|---|
| AsegOR13_F | CGGGATCCGCCACCATGAATTCGATTCTTAAAACTTTGGAAGACC |
| AsegOR13_R | GCTCTAGATTACTTTTTCAGCAATGTGTAGTAACTGTAC |
| AsegOR14_F | CGGGATCCGCCACCATGGAAATCAAAACGTCCTCAGAAGT |
| AsegOR14_R | GCTCTAGATTAAGCTGTGATCAAATTAAAGAAGGAAAAAGAA |
| AsegOR15_F | GAAGGCCTGCCACC ATGATTCAAGCATACAAAACGTACGATG |
| AsegOR15_R | GCTCTAGATTATTCTGAATATGCAGTACGAAGCAAATTG |
| AsegOR16_F | CGGGATCCGCCACCATGGATGAAGAAGTAGAATTCAAGCCAT |
| AsegOR16_R | GCTCTAGATTATGACCTCAATAGCGTAAACATTGAATATG |
| AsegOR17_F | CGGGATCCGCCACCATGATAACCTTACGAGAAATTGAACATGAAATC |
| AsegOR17_R | GCTCTAGACTACTTCATAGTAGACCGAAGAGCAGTAT |
| AsegOR18_F | CGGAATTCGCCACCATGCGTCTTCAAATAATAAAAGATTTTCTCTTGA |
| AsegOR18_R | GCTCTAGATTATTTGCTGTATTGACGTAGCATATTGAAATAT |
| AsegOR19_F | CGGGATCCGCCACCATGGAATTGCATCAAATAGACTGCTTTAAAATA |
| AsegOR19_R | GCTCTAGATCATACTGATTTAGCTCTTAATAAAACAAAAAGATCGT |
| AsegOR20_F | CGGGATCCGCCACCATGGAAGAAAACCCACTTCTCATTGA |
| AsegOR20_R | GCTCTAGATTAAGTTGGTGAGCTGTATAATGTTTGC |
